# Quantitative Mapping of Organelle Positioning in Cultured Cells Using Semi-Automated Image Analysis Pipeline

**DOI:** 10.64898/2026.04.24.720625

**Authors:** Katerina Jerabkova-Roda, Vincent Hyenne, Jacky G. Goetz

**Affiliations:** Tumor Biomechanics, Strasbourg, France; INSERM UMR_S1109, Strasbourg, France; Université de Strasbourg, Strasbourg, France; Fédération de Médecine Translationnelle de Strasbourg (FMTS), Strasbourg, France; Equipe Labellisée Ligue Contre le Cancer; CNRS, SNC5055, Strasbourg, France

**Keywords:** Subcellular architecture, Organelle positioning, Live-cell imaging, Immunofluorescence, Lysosomes, Radial distribution, CellProfiler, ImageJ

## Abstract

Subcellular architecture is tightly controlled and contributes to the maintenance of cell’s homeostasis. Organelles are regulated in size, shape, number and position which respond to changes in extracellular environment. Lysosomes are of particular interest as they integrate various functions in the cells (nutrient sensing, metabolism, cell migration and adhesion), serving as signaling hubs. Their function is tightly linked to their subcellular position and deregulation of lysosome homeostasis leads to several diseases including cancer. Therefore, methods allowing precise analysis of organelle subcellular distribution can aid in fundamental, diagnostic and therapeutic approaches. Here, we provide a versatile image analysis pipeline using ImageJ and CellProfiler. This workflow allows to quantify subcellular lysosome distribution in living and fixed melanoma cells, and is applicable to other subcellular compartments and to various cell types.

## 1. Introduction

Melanoma progression is accompanied by changes in their transcription program leading to expression of matrix-degrading enzymes and to invasion and progression into metastatic stages ***(1)***. Large efforts were made in studying the transcriptional programs of melanoma cells, yet their functional implications on the cellular level remain elusive. Tight regulation of subcellular architecture is important for maintenance of cell’s homeostasis. Organelles often mediate intracellular signaling events and, based on their morphology, position within the cell, and contacts with other organelles, they exert different biological functions ***(2)***. Such plasticity, although essential, also allows cancer cells to quickly adapt to the changing environment. The Golgi apparatus is responsible for protein processing and trafficking, and secretion of proteins forming the extracellular matrix. In cancer, its architecture oscillates between compacted and vesicular, fragmented states, which allows it to modify the tumor microenvironment and thus stimulate cancer progression via altered protein secretion of cytokines, proteases, proteoglycans and procollagen ***(3)***. Also, mitochondrial morphology has direct impact on tumor growth and metastasis. Many RAS-driven cancers have fragmented mitochondria, including pancreatic ***(4)***, lung ***(5)***, liver ***(6)*** and squamous cell carcinoma ***(7)***, In contrast, mitochondria fusion factor Opa1, is essential for tumor cell survival in lung adenocarcinoma ***(5)***. Furthermore, inter-organelle contacts directly impact the subcellular architecture as lysosome-mitochondria membrane contacts can mediate mitochondria fission ***(8)*** and contacts of lysosomes with the endoplasmic reticulum (ER) can modulate the lysosome motility ***(9)***. The endo-lysosomal system has various functions, including degradation, metabolic signaling, gene regulation, plasma membrane repair, and cell adhesion and migration ***(10)***. In response to the environment, lysosomes are able to change their position, size, number, composition and function ***(11–13)*** which serves as a coping mechanism to adapt to homeostatic deviations such as stress (nutritional, UV, osmotic) and mechanical forces ***(14, 15)***. Lysosomes are transported to cell periphery along microtubules via kinesins ***(16)***, which leads to lysosome exocytosis and contributes to cancer cell invasion ***(12)*** and chemoresistance ***(17)***, (Fig. 1A).

**Figure 1.**
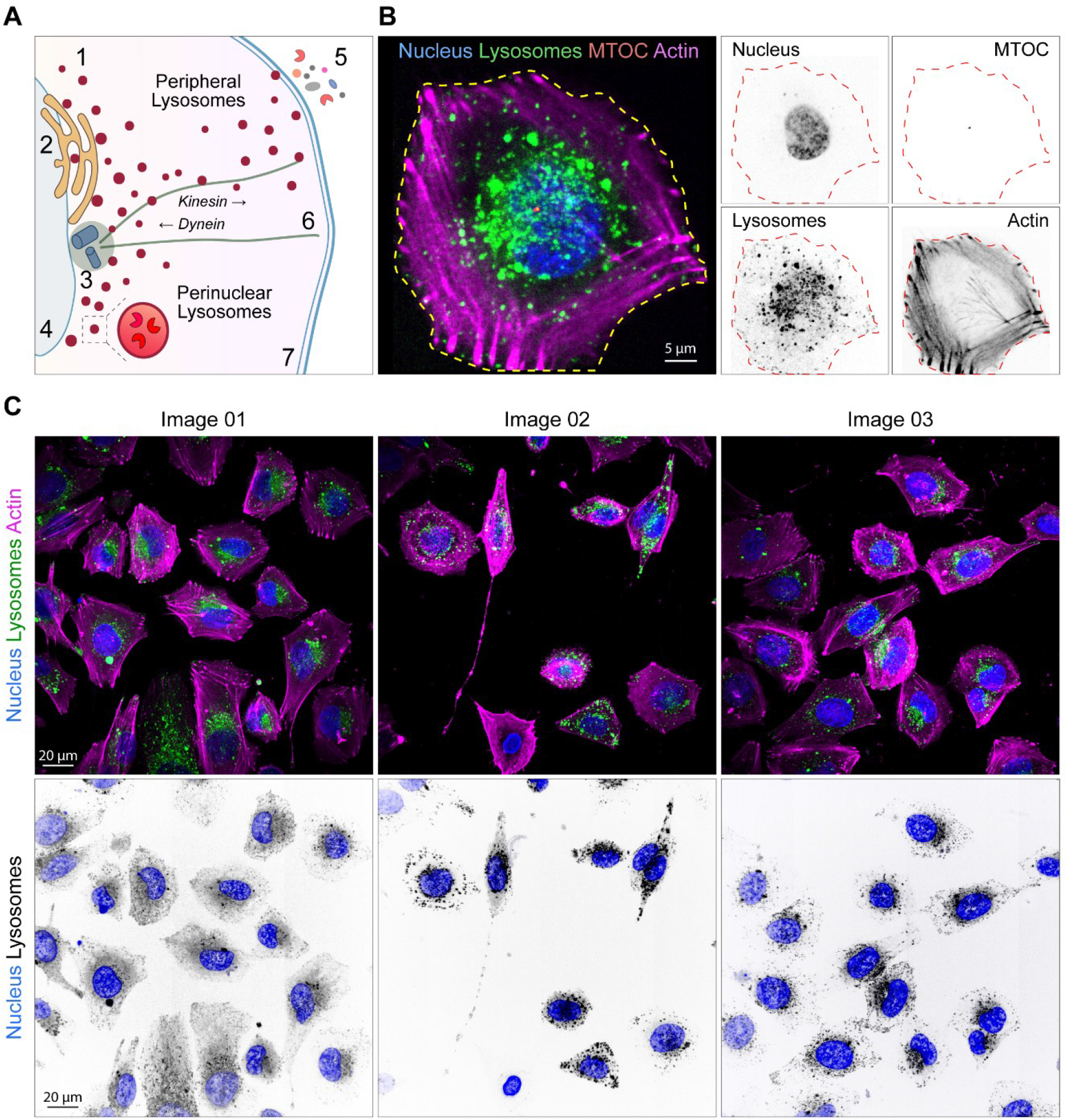
Lysosome positioning in cultured melanoma cells. A) Illustration of lysosomal dynamics in living cells. 1. Lysosomes, 2. Endoplasmic reticulum, 3. Centrosome, 4. Nucleus, 5. Secreted factors, lysosomal exocytosis, 6. Microtubules, 7. Plasma membrane. B-C) Live-cell imaging of WM983B melanoma cells by spinning disk microscopy, maximum intensity projection. Cells are stained for lysosomes (Lysotracker green), nucleus (NucBlue), actin (SiR-Actin647, magenta), and microtubule organizing center (stable expression of cenexin-tdTomato). B) Left: Representative image of a single cell, yellow line= cell border, drawn manually. Right: Single channels showing individual markers. C) Three different example images showing stained melanoma cells. Top: Composite image, merge of lysosomes (green), nucleus (blue) and actin (magenta). Bottom: Inverted fluorescent images showing nucleus (blue) and lysosomes (grays).

To understand how these processes impact cancer biology and disease outcome, it is necessary to develop methods allowing to quickly and reliably assess the organelle subcellular distribution as it can provide a valuable insight on mechanisms of cancer progression and diagnosis. This method was optimized for lysosome positioning analysis in melanoma cells, but is applicable to any subcellular compartment marker which can be reliably labelled by live cell imaging or by immunofluorescence. To date, several approaches to quantitatively assess lysosome positioning have been developed. These include: i) Manual scoring in a blinded setup ***(18)***, ii) Lysosome segmentation followed by distance measurements to the nucleus ***(19)***, to the cell periphery ***(20)***, to the microtubule organizing center (MTOC) ***(21)***, to the nearest neighbor ***(12, 22)***, iii) Segmentation and object counting near the nucleus and in the cell periphery ***(17)***, iv) Intensity-based quantifications, calculating the perinuclear index ***(23, 24)***, the mean distribution from edge ***(25)*** and the radial distribution between cell periphery and the nucleus ***(12)*** or the centrosome ***(15)***. These methods often use software that is not available free of charge and it commonly requires the use of a custom-written code. Many of these methods rely on lysosome segmentation which can be difficult, especially when lysosomes are clustered around the nucleus. Therefore, intensity-measurements represent more versatile approach to assess their subcellular distribution.

Here, we provide a robust, intensity-based analysis pipeline, using two free, open-source softwares, CellProfiler ***(26)*** and ImageJ/Fiji ***(27)***, together with a detailed step by step protocol. This protocol first describes the cell culture, cell staining and imaging, followed by a three-step analysis pipeline: i) cell and nucleus segmentation in ImageJ / Fiji, ii) lysosome radial distribution measurements in CellProfiler and iii) data visualization strategies.

## 2. Materials

### 2.1. Cell culture, cell staining and imaging

1. Cell line: WM983B, human metastatic melanoma cell line mutant for BRAF^V600E^ (*see* **Note 1**).
2. TSM medium, MCDB153 and Leibovitz’s L-15 medium in a 4 to 1 ratio, respectively, supplemented with 2% fetal bovine serum, 1.68 mM CaCl_2_ and 1% penicillin/streptomycin (stock 10.000 U/ml Penicillin, 10 mg/ml Streptomycin).
3. Trypsin-EDTA solution, 0.05% trypsin
4. optional: EDTA (Versene) 1% in PBS, free from Ca^2+^ and Mg^2+^
5. 25 cm^2^ flasks with filter cup, Glass-bottom dishes suitable for confocal imaging with #1.5 coverslip thickness (4-chamber, 35mm, glass bottom dish)
6. Fibronectin coating solution, 10μg/ml fibronectin in water or in PBS
7. Trypan blue stain, 0.4% solution to distinguish viable cells during counting
8. optional: Countess™ 3 Automated Cell Counter
9. **Actin staining solution (1x)**, volume for 4 wells: 1.3 ml TSM medium + 1.3 μl SiR-actin probe (1:1000 dilution, without Verapamil). **Lysosome staining solution (2x)**, volume for 4 wells: 1.25 ml TSM medium + 1 drop NucBlue Live Probe + 2.5 μl Lysotracker green (from 50 μM stock, final staining concentration 50 nM), (*see* **Note 2**).
10. Microscopy: Inverted Spinning Disk microscope (60x objective NA 1.2 or similar), equipped with humidified chamber keeping the temperature at 37°C with 5% CO_2_ atmosphere (essential for live-cell imaging).

### 2.2. Analysis pipeline - Software

1. ImageJ / Fiji (version 1.54p, Java 21.0.7) ***(27)***
2. biovoxxel plugin for Fiji, https://doi.org/10.5281/zenodo.5987322. Downloaded from www.biovoxxel.de/tools/ using the configuration file ‘BioVoxxel_Fiji_Config.groovy’ ***(28)***
3. CellProfiler (version 4.2.1) ***(26)***
4. GraphPad Prism, version 11.0.0 (84) for Windows 64-bit, GraphPad Software, Boston, Massachusetts USA, www.graphpad.com
5. Spreadsheet editor (Microsoft Excel or similar)

## 3. Methods

### 3.1. Cell culture, cell staining and imaging

1. Culture the cells in 5% CO_2_ incubator at 37 °C using TSM medium and passage them 1-2 times a week in a 1:5 dilution in 25 cm^2^ flasks with filter cup.
2. Coat the 4-compartment glass bottom dish with fibronectin (10 μg/ml), 120 μl / coverslip, incubate at 37 °C for at least one hour before cell seeding. Wash the dish one time with 500 μl of warm culture medium.
3. Wash cells two times with 2 ml of warm PBS, remove PBS and add trypsin (500 μl / T25 flask), incubate 1 min at 37 °C and resuspend thoroughly in warm culture medium (5ml / T25 flask) to achieve suspension of single cells (*see* **Note 3**).
4. Day 0: Count cells (use Trypan Blue to count the living cells, 1:1 ratio), seed 50 000 cells per compartment, stain and image the cells on the next day, Day 1 (*see* **Note 4**). Cells should be at 50 – 80% confluency at the time of imaging.
5. Day 1: Stain the cells with Actin staining solution, 1.3 ml per 4-chamber dish in total. First, remove culture medium, then add 300 μl actin staining solution per well and incubate 45 minutes at 37 °C (*see* **Note 5**).
6. Pre-heat the microscope to stabilize the imaging chamber at 37 °C with 5% CO_2_ prior to imaging.
7. To stain nuclei and lysosomes, prepare Lysosome staining solution (2x), mix and add 300 μl to each well (total volume is 600 μl / well), incubate at 37 °C for additional 15-20 minutes. Cells are ready to be imaged (*see* **Note 6**).
8. Microscopy: Image cells, stained with selected markers (Fig. 1B), choose regions where cells are not confluent (Fig. 1C), isolated cells will be easier to segment. Acquire z-stacks of the full cell volume (z-step 0.2 μm - 0.5 μm), image at least 30 cells / condition (around five fields of view), keep the same z-step and number of slices, and constant laser power and exposure time in all the images.

### 3.2. Analysis pipeline – Step 1: Segmentation

First, segment cells and nuclei in Fiji (*see* **Note 7**). This macro saves maximum projections for each channel, generates binary masks for cells and nuclei and saves the ROIs of all segmented objects. The “Segmentation_LTrace.ijm” macro is available in Zenodo archive ***(29)***.

1. Image pre-processing: prepare input folder with multi-channel z-stacks (3- or 4-channel images are supported), saved in a TIF format.
2. Run the “Segmentation_LTrace” macro in Fiji and select an input folder, all images in this folder will be analyzed.
3. Select the output folder, all images will be saved there. This folder will be used as input for the next analysis step in CellProfiler.
4. Assign channels to markers for nucleus (or MTOC), target compartment (lysosomes) and cell border (actin). This macro works with three- and four-channel images (the fourth channel is not analyzed).
5. Nucleus segmentation: Adjust the threshold to correctly segment all nuclei (holes will be filled in the next step using ‘Fill Holes’ command and small particles will be filtered out using ‘Analyze Particles’ command, to keep only the nuclei).
6. Cell segmentation method is based on thresholding and marker-controlled watershed (available as part of the biovoxxel plugin in FiJi). This macro was designed to enhance even faintly visible structures using Enhance Local Contrast (CLAHE) command (*see* **Note 8**). Follow these steps as they appear in the macro:
  a. Adjust threshold (Li) to best capture the fine outlines of cells (holes will be filled in later steps), then adjust threshold (MinError) to have the cell area covered (eliminate holes, cells can be touching) (Fig. 2A).
  b. Cell edges are automatically created using the ‘Find Edges’ command (Fig. 2B).
  c. Identify maxima (Fig. 2C). **Important**: Select ‘Maxima Within Tolerance’ from the drop-down menu. Using ‘Preview’, select the highest prominence number that allows to identify the individual cells. If cells are over-segmented, the prominence value is too low.
  d. Marker controlled watershed uses three images: binary cell mask (Li Threshold), Edged image (Find Edges) and Label (from Find Maxima step in Distance Map) (Fig. 2A, B, D). These images are automatically loaded to create the watershed image.
  e. Refine segmentation – these two steps allow for correction of segmentation errors in the original Watershed image (Fig. 2E left). Individual objects are labeled by different colors. First, with brush tool (size 2pt) draw black separating lines where two cells need to be distinguished. Second, draw white lines where you want to connect two objects (Fig. 2E right).
  f. Select cells that will be analyzed for lysosome positioning. Only whole cells can be analyzed, cells spanning outside the image border should be excluded. Click on each cell with ‘wand’ tool and press ‘t’ to add this selection to the ROI manager. Repeat this step for all the cells that you wish to analyze (Fig. 2F).
7. The Fiji output folder now contains three subfolders: i) ‘Fluo_images_input’ (containing maximum intensity projections for each channel, used in Step 2), ii) ‘Masks_input’ (containing binary masks for segmented cells and nuclei (Fig. 2G), used in Step 2), and ‘Other_files’ which contains the ROIs of segmented objects and other images that can be used for the watershed (Edges, Label, Li threshold) or for quality control and adjustments of the segmentation (Refined binary mask, Watershed image).

**Figure 2:**
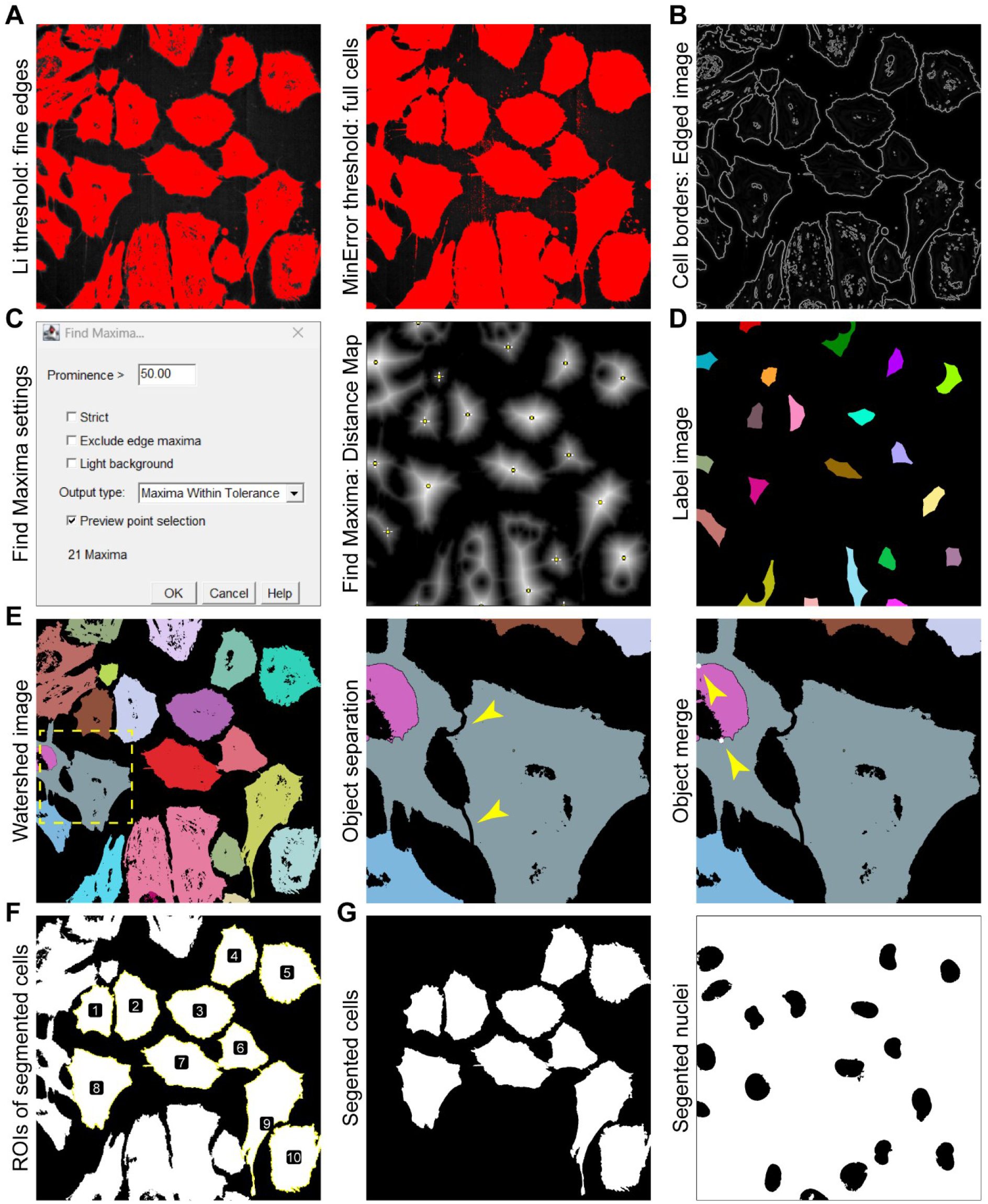
Overview of analysis pipeline for Fiji (Step 1). A) Thresholding methods for cell segmentation. B) Image with cell outlines, result of Find Edges command. C) Settings used to identify local maxima and thus to distinguish the individual cells. D) Label image calculated from the Local Maxima. E) Marker-controlled watershed image. Zoomed regions shows examples of object separation (left) and object merge (right). F) Selected cells and their ROIs, after manual selection step. G) Binary masks of segmented cells and segmented nuclei.

### 3.3. Analysis pipeline – Step 2: CellProfiler analysis

1. Open the analysis pipeline “LTrace_RadialDistribution” in Cell Profiler. The pipeline is available in Zenodo archive ***(29)*** and its overview is illustrated in Fig. 3A.
2. Drag and drop the Fiji output folder to the section called ‘Images’ and update the ‘NamesAndTypes’ tab (Fig. 3B). Images should be automatically grouped, verify that labels are correctly assigned to individual image sets (*see* **Note 9**).
3. Define CellProfiler output folder in File → Preferences → Default output folder (select a path). All analysis files will be saved here.
4. The analysis pipeline generates cell and nucleus objects from the binary masks (Fiji output) and selects cells containing single nucleus for Radial distribution analysis of lysosome signal (Fig. 3C), (*see* **Note 10**).
5. The output folder contains:
  a. Images: i) cell and nuclei overlay images annotated with object numbers for each selected object (Fig. 3D). Important: Nuclei do not always have the same object number as the cells. ii) Radial distribution maps, which represent the lysosome subcellular distribution. Each cell is divided into 9 concentric segments (Bins), while bin 1 lies close to nucleus and bin 9 close to plasma membrane. Mean lysosome intensity is measured in each bin and is indicative of lysosome localization pattern. High mean intensity value (high lysosome density) is represented by dark red color and areas (bins) with fewer lysosomes are marked by lighter colors (Fig. 3E).
  b. Measurements: i) ‘Cells_single_nuc’ (This file contains measurements for the selected cells: Image and object number, file name, cell area in px= AreaShape_Area, Integrated and Mean lysosome intensity per cell, Radial distribution= MeanFrac_Lysosomes bin 1 – bin 9). ii) ‘SelectedNuclei’ (This file contains measurements for the selected nuclei: Image and object number, file name, nucleus area in px= AreaShape_Area and several shape parameters (Compactness, Eccentricity, Form Factor, perimeter, major and minor axis length). iii) Other files that are not needed for this analysis.

**Figure 3:**
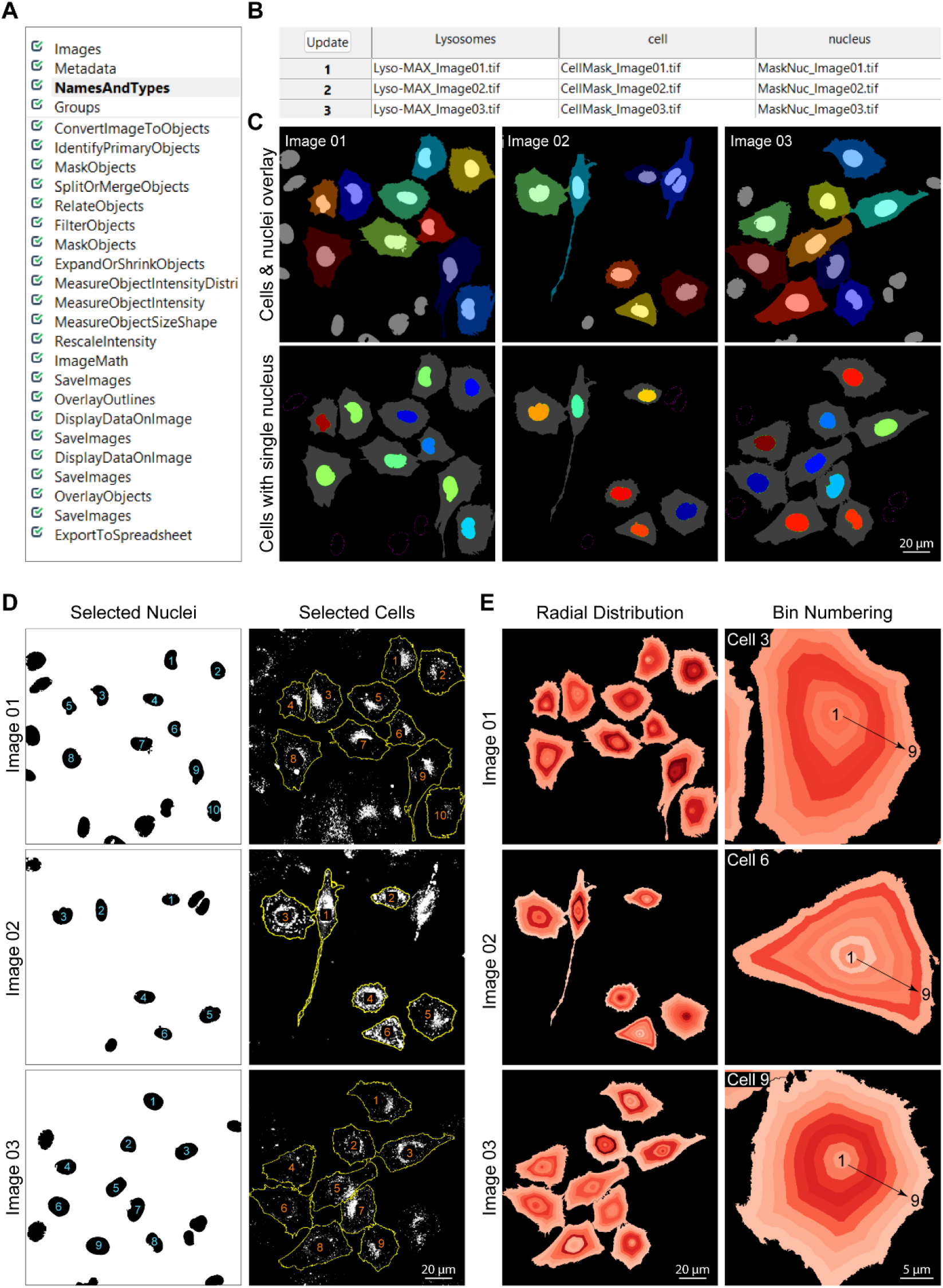
Overview of the CellProfiler analysis pipeline (Step 2). A) Analysis pipeline, list of all steps. B) Names assigned to individual images. C) Selection of cells with single nuclei for the radial distribution analysis. D) Examples of CellProfiler output files. Left: Overlay image shows all segmented nuclei, numbers annotate only nuclei of analyzed cells. Right: Analyzed cells are numbered and their outlines are overlaid on lysosome image. E) Radial distribution map is color-coded based on lysosome intensity. Dark red = high intensity. Right panel shows zoomed regions, 1 cell per image, showing the concentric bins, numbered from 1 (central) to 9 (peripheral).

### 3.4. Analysis pipeline – Step 3: Data visualization

1. Work with file called ‘Cells_single_nuc’ and focus on values ‘MeanFrac_Lysosomes’ for bin 1 – bin 9. These correspond to the Radial distribution measurement. Each cell is divided into nine concentric bins, mean lysosome intensity is provided for each bin (bin 1 = perinuclear, bin 9 = peripheral). To visualize lysosome subcellular distribution, normalize the intensity of each bin to the average intensity of bins 1-9 in each cell. Example quantification and test images are available as Supplementary Material.
2. Plot the normalized values in GraphPad Prism using xy plot to visualize the lysosome distribution. As an example, we first show intensity profiles for three selected cells (Fig. 4A, B). Simplified view can be achieved by plotting Area Under the Curve (AUC) values for the three cell regions: i) Perinuclear (bin 1-3), ii) Mid (bin 4-6), iii) Peripheral (bin 7-9), Fig. 4C. We can further visualize the experimental mean for each condition by plotting replicate values (Fig. 4D) or the individual values for each cell (Fig. 4E).
3. For quantitative assessment, plot the AUC values for individual regions: Perinuclear (bin 1-3), Mid (bin 4-6), Peripheral (bin 7-9). AUC analysis is available in GraphPad Prism. Identify the smallest value in the dataset and set it as baseline for all AUC calculations (y= MIN), plot the AUC values for each region (Fig 4F). Example quantification and test images are available as Supplementary Material.

**Figure 4:**
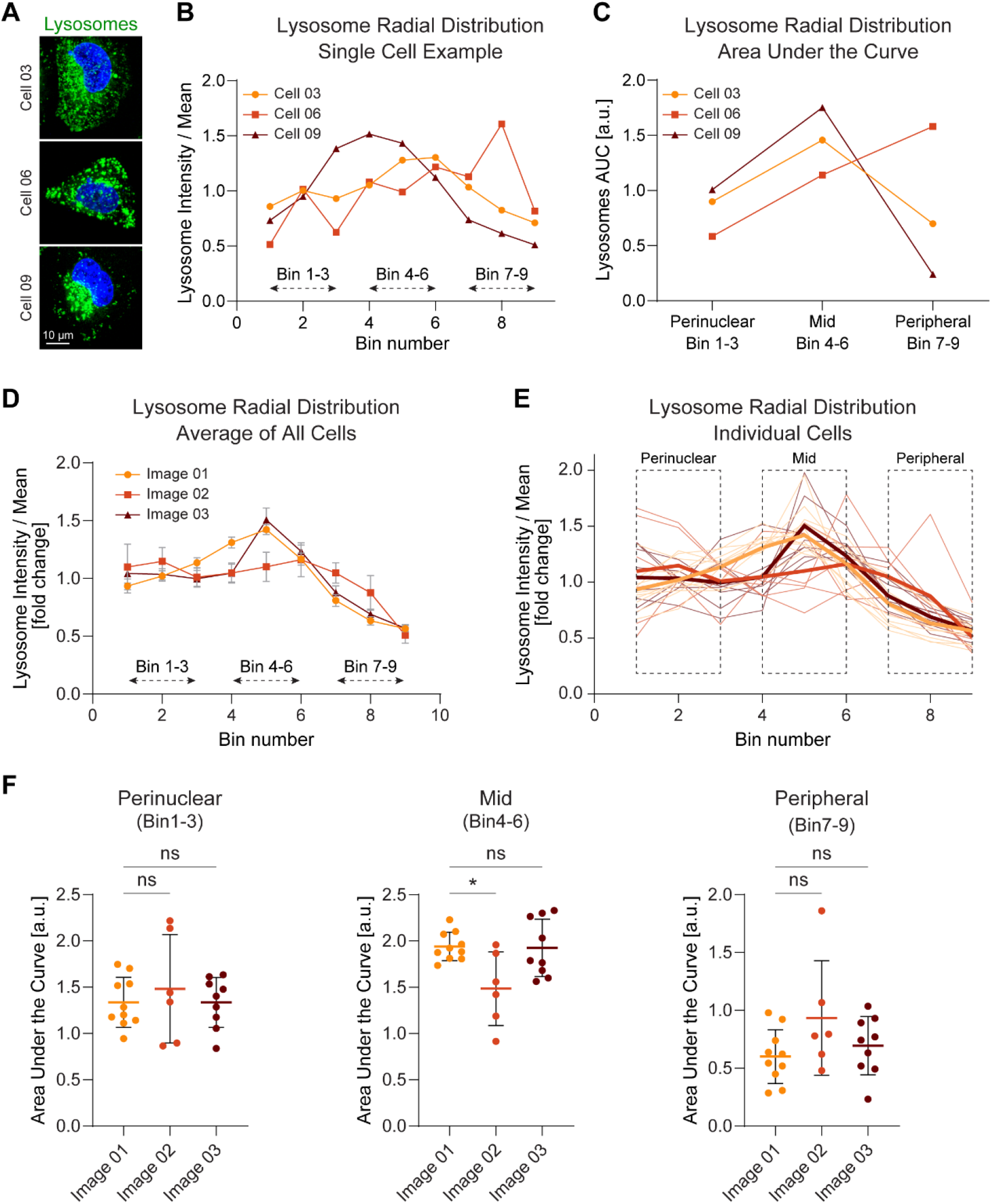
Quantitative representation of lysosome subcellular distribution (Step 3). A) Fluorescent images, single cell examples, lysosomes (green), nuclei (blue). B) Intensity profile for three individual cells shown as normalized radial distribution: Mean intensity in each bin, normalized to the mean intensity per cell. C) Area under the curve for each cell in three different cell regions: Perinuclear, Mid, Peripheral. D-E) Quantification of lysosome distribution. Image 01: n= 10 cells, Image 02: n= 6 cells, Image 03: n= 9 cells. Normalized radial distribution, shown as D) Mean value per image and E) Value per individual cells. Thick line shows the mean value per experiment. F) Area under the curve quantification in the three cell regions: Perinuclear, Mid, Peripheral as quantified in GraphPad Prism 9. Statistics: Kruskal-Wallis with Dunn’s multiple comparisons test.

## Supporting information

Example quantification of normalized radial distribution

## 4. Notes

**Note 1** The analysis was optimized using a patient-derived melanoma cell line, but the protocol is suitable for any adherent cells, cancerous or non-cancerous.

**Note 2** To delimit the cell borders, other dyes can be used: SiR-Tubulin (microtubules), membrane dyes (i.e. MemGlow, CellMask, Wheat Germ Agglutinin (WGA-AlexaFluor)) or dyes staining cytoplasm (i.e. CellTrace), to name just a few. In addition, the quantification can be done on fixed samples, stained by immunofluorescence, using markers for the selected compartments (i.e. LAMP1 for lysosomes, TOM20 for mitochondria, calnexin for endoplasmic reticulum, and cenexin / γ-tubulin for MTOC).

**Note 3** Cells can be washed with 1% EDTA (fast wash, or 1-minute incubation at 37°C) prior to incubation with trypsin to better dissociate the cells and achieve single cell suspension.

**Note 4** For experiments requiring gene knockdown by siRNA, seed 20 000 – 30 000 cells per well and follow the Lipofectamine RNAiMAX forward transfection protocol (24 – 48 hours of siRNA treatment prior the cell staining step). If reproducible cell architecture is required, cells can be also seeded on micropatterns.

**Note 5** Alternatives: i) Cells stably expressing fluorescently tagged marker proteins can be used for live-cell imaging, ii) Immunofluorescent labelling of selected markers can be done on fixed cells, as explained in Note 2.

**Note 6** To minimize autofluorescence, use cell culture medium without phenol red.

**Note 7** This step can be replaced by alternative methods for cell and nucleus segmentation that produce binary masks. Pay attention to the file names to ensure that images are properly recognized in CellProfiler. Image containing segmented cells should start with “CellMask_” and image of segmented nuclei should start with “MaskNuc_”.

**Note 8** If your cells are labelled by bright cytoplasmic marker, you can skip this step by removing lines 171 and 180 from the macro.

**Note 9** To minimize errors, use two- or three-digit numbers (Name_01 or Name_001) and avoid using spaces in the file names.

**Note 10** This pipeline analyses radial distribution around centers of nuclei. Distribution around MTOC (microtubule organizing center) is more accurate as MTOC is not always lying at the center of nucleus, but often on the side (Fig. 1B). Segmented centrosomes can be used instead of segmented nuclei.

## Data availability

The ImageJ macro, the CellProfiler pipeline and several example images and analysis file are available on Github: github.com/KJerabkovaRoda/LTrace. The ImageJ macro and the CellProfiler pipeline were deposited to the Zenodo archive: doi.org/10.5281/zenodo.19334736 ***(29)***. Example images are available as Supplementary Material.

## Acknowledgements

The work has been initially supported by NANOTUMOR Consortium, a program from ITMO Cancer of AVIESAN (Alliance Nationale pour les Sciences de la Vie et de la Santé, National Alliance for Life Sciences &Health) within the framework of the Cancer Plan (France). It was later supported – in the Tumor Biomechanics Lab - by INCa (Institut National Du Cancer), La Ligue contre le Cancer, ARC (Association pour la Recherche contre le Cancer), FRM (Fondation pour la Recherche Médicale), the National Plan Cancer initiative, the Region Est, INSERM, University of Strasbourg, RAS Foundation, TDLR (Trailers de la Rose), Club Féminin Lampertheim & Rohan Athlétisme Saverne. We are grateful to PIC-STRA imaging platform (CRBS, Strasbourg: P. Kessler) and members of the Tumor Biomechanics lab for their help. We acknowledge the use of ChatGPT (OpenAI, GPT-5.3; Bio-Image Analysis GPT by Robert Haase) during the initial design to decide which steps to include in the ImageJ analysis workflow. All implemented suggestions were thoroughly analyzed, validated, and verified by the authors. We also thank ChatGPT for assistance with drafting Alt text for figures. All output was carefully reviewed and revised by the authors. We thank M. Théry (CytoMorpho Lab, Grenoble) for providing the cenexin construct to label MTOC in living cells.

